# Mitigation of imprinted antibody responses against COVID-19 in highly vaccinated older adults

**DOI:** 10.64898/2026.05.21.725708

**Authors:** Rebecca B. Morse, Daniel J.S. Egan, Mark Tsz Kin Cheng, Mazharul Altaf, Kimia Kamelian, Claire Gallagher, Lourdes Ceron Gutierrez, Olga Sokolova, John R. Bradley, Kenneth G.C. Smith, Sarah Meloy, Gillian Ison, Anita Furlong, Ingrid Scholtes, Emma Le Gresley, Rainer Doffinger, Chee Wah Tan, Ravindra K. Gupta

**Affiliations:** Department of Medicine, University of Cambridge, Cambridge, UK; Cambridge Institute of Therapeutic Immunology & Infectious Disease (CITIID), Cambridge, UK; Addenbrooke’s Hospital, Cambridge University Hospitals NHS Foundation Trust, Cambridge, UK; The Walter and Eliza Hall Institute of Medical Research (WEHI), Parkville, VIC, Australia; Department of Medical Biology, University of Melbourne, Melbourne, VIC, Australia; Program in Emerging Infectious Diseases, Duke-National University of Singapore Medical School, Singapore; Infectious Diseases Translational Research Programme, Department of Microbiology and Immunology, Yong Loo Lin School of Medicine, National University of Singapore, Singapore; HKJC Global Health Institute, Hong Kong, SAR China; Africa Health Research Institute, Durban, KwaZulu-Natal, South Africa

**Keywords:** SARS-CoV-2, COVID-19, Immune Imprinting, Vaccination, Antibodies, Elderly

## Abstract

SARS-CoV-2 continues to evolve from the Omicron serotype, with BA.2.86 sublineage JN.1 and descendants predominating in 2025-26 and recent emergence of the highly divergent BA.3.2 saltation variant. Elderly individuals continue to be at greatest risk of clinical complications from COVID-19, yet contemporary data on kinetics of immune potency and breadth following multiple vaccinations remain limited in this group. We studied a cohort of forty-three healthy older adults (median age = 85 years, IQR 75-88, 40% female). Using both pseudotyped virus (PVNT) and surrogate virus (SVNT) neutralization-based assays, we demonstrate that JN.1 and KP.2 vaccinations six months apart elicit high potency neutralization across all studied variants except BA.3.2.2, where titres were around 2-3 fold lower. Waning of serum neutralizing activity was modest between vaccine doses, suggesting sustained immunity following multiple vaccines. Importantly, the half-life of serum neutralization against Wu-1 (179 days) was substantially longer compared to JN.1 subvariants (43.7-70.1days), consistent with imprinted and longer-lived Wu-1-specific responses. Furthermore, half-life for BA.3.2.2 neutralisation was prolonged at 107 days compared to JN.1 subvariants, implicating imprinted ancestral variant targeting antibodies in neutralization of this saltation variant. While absolute neutralization titers remained highest against ancestral Wu-1 at all timepoints due to multiple historical exposures and accumulation, the recall responses after the recent KP.2 vaccine revealed a shift in immunodominance: neutralization against full-length Wu-1 spike was not boosted, whereas all tested JN.1 descendants and BA.3.2.2 showed significant boosts, indicating that immune imprinting against ancestral Wu-1 was partially overcome. These data indicate that in the elderly, protective neutralizing antibody responses against recent VOC can be achieved, with alleviation of immune imprinting and reduction of waned responses through biannual, strain-updated booster vaccines. Imprinted antibodies against ancestral pre Omicron variants may still be relevant for protection against further highly immune evasive variants such as BA.3.2.2, highlighting that careful characterisation of population level COVID-19 humoral immunity is still warranted.

## Introduction

The emergence of severe acute respiratory syndrome virus 2 (SARS-CoV-2) in late 2019 led to the rapid development and deployment of vaccines to protect against infection and severe coronavirus disease (COVID-19). While first-generation vaccines targeting the original Wuhan-Hu-1 (Wu-1) strain were effective in most individuals, viral evolution towards increased immune evasion and transmissibility led to widespread reinfections. This diversification is thought to be driven in part by intrahost evolution during chronic infections (1,2). With the emergence of the Omicron variant and its sub-lineages presenting a marked change in the spike protein (3,4), immune-evasive variants continue to appear periodically with a subsequent decrease in Wu-1-based vaccine efficacy, prompting production of seasonal, variant-specific ‘booster’ vaccines. Recommendations first transitioned to incorporate Omicron in bivalent formulations targeting BA.1 or BA.4/5(5–12) and increased antibody titers against Omicron were observed, but breadth was similar in people receiving monovalent Original Wu-1 vaccines (9,13,14). The preferential recall of B cell responses against conserved epitopes amongst variants was observed, supporting immune imprinting against the Original Wu-1 strain(15–17). However, these epitopes were often non-neutralizing, and recommendations therefore shifted to promoting monovalent Omicron variant vaccines in the hopes of reducing ‘backboosting’. Some studies have shown mitigation of immune imprinting and *de novo* responses via repeated Omicron exposures, whereas others have confirmed continued recall of shared epitopes with Wu-1. Nevertheless, serum neutralizing titers against Wu-1 spike continue to be higher than against Omicron in almost all cohorts(18–21).

Following the dominance of the BA.2.86 sublineage JN.1 in 2024, the contemporary antigenic landscape shifted towards its derivatives such as NB.1.8.1 and XFG. Concurrently, the highly divergent BA.3.2 saltation variant and its sublineages have emerged as a modest but persistent fraction of reported cases(22). Within the UK and Europe, there has been steady growth of BA.3.2.2 in recent months, with an increased proportion of reported cases amongst hospitalized children(23). The WHO Technical Advisory Group on COVID-19 Vaccine Composition (TAG-CO-VAC) periodically requests data to monitor decreased susceptibility to neutralization and ongoing accumulation of mutations—for example, to understand immunity against more recent NB.1.8.1, XFG and BA.3.2 lineages designated of interest—thereby informing COVID-19 vaccine composition(24).

As older adults and immunocompromised individuals continue to experience the highest rates of hospitalization and death due to COVID-19, these groups have been prioritized for repeated seasonal vaccinations(25,26). Older adults exhibit distinct humoral challenges, including lower peak antibody titers, reduced neutralization breadth and antibody-dependent cellular cytotoxicity (ADCC) responses, atypical B cell populations, and accelerated antibody titer decay compared to younger individuals following COVID-19 vaccination(27–32), although this effect is diminished by multiple booster vaccines(33–35). in Current UK COVID-19 vaccination campaign offers biannual ‘booster’ vaccines and is targeted towards all adults >75 years old and immunosuppressed individuals >6 months old(26). These ‘boosters’ consist of WHO-approved variant-specific monovalent vaccines evaluated by the TAG-CO-VAC(24,26) and developed to bolster immunity to circulating Omicron variants that evade immunity established against the original Wu-1 strain(36–38). However, few studies have measured waning after sequential vaccinations in the JN.1 lineage era and little information exists about older adults. Thus, our understanding of the durability and breadth of immunity elicited by contemporary boosters in the elderly remains limited despite this population being the main recipients of these vaccines and the rapid ongoing emergence of new variants evading immunity.

Here we report on the humoral immunity to globally relevant JN.1 subvariants and the BA.3.2.2 variant within a highly vaccinated UK elderly cohort. Using a full-spike pseudotyped virus neutralization assay (PVNT) along with an RBD-based 21-plex surrogate virus neutralization test (SVNT), we characterized both full-spike and domain-specific neutralizing antibody titers and breadth. We deployed these results to construct antigenic maps across timepoints to describe the shift in antigenic landscape following boosting with the recent KP.2 mRNA vaccine. These data contribute important insights into how updated monovalent vaccines modulate immune imprinting and breadth of immune responses in a clinically relevant population facing an ongoing wave of highly divergent SARS-CoV-2 viral evolution.

## Results

We studied a cohort of forty-three healthy older adults (median age = 85 years, IQR 75-88, 40% female) who received a COVID-19 vaccine during the UK Autumn 2025 vaccination campaign (Table 1). These participants were all recently sampled following administration of Comirnaty KP.2 (n = 41) or Comirnaty LP.8.1 (n = 2) variant monovalent mRNA vaccine during the Autumn 2025 vaccination window (Figure 1A, Supplementary Figure 1A). The median time between the autumn 2025 dose and previous dose was 182 days (IQR 174-190). Consistent with UK National Health Service guidelines to be vaccinated biannually, most (n = 39/43, 90.7%) individuals received a JN.1 vaccine during both the autumn 2024 and spring 2025 vaccine campaigns, followed by a KP.2 vaccine in the autumn 2025 campaign (Supplementary Figure 1A). One individual received a JN.1 dose in autumn 2024, but a KP.2 dose in spring 2025 and an LP.8.1 dose in autumn 2025. Three individuals only received annual doses after the primary series (seven doses including the autumn 2025 campaign), with all receiving a JN.1 vaccine in autumn 2024 followed by a KP.2 (n = 2) or LP.8.1 (n = 1) vaccine in autumn 2025 (Supplementary Figure 1A).

**Figure 1.**
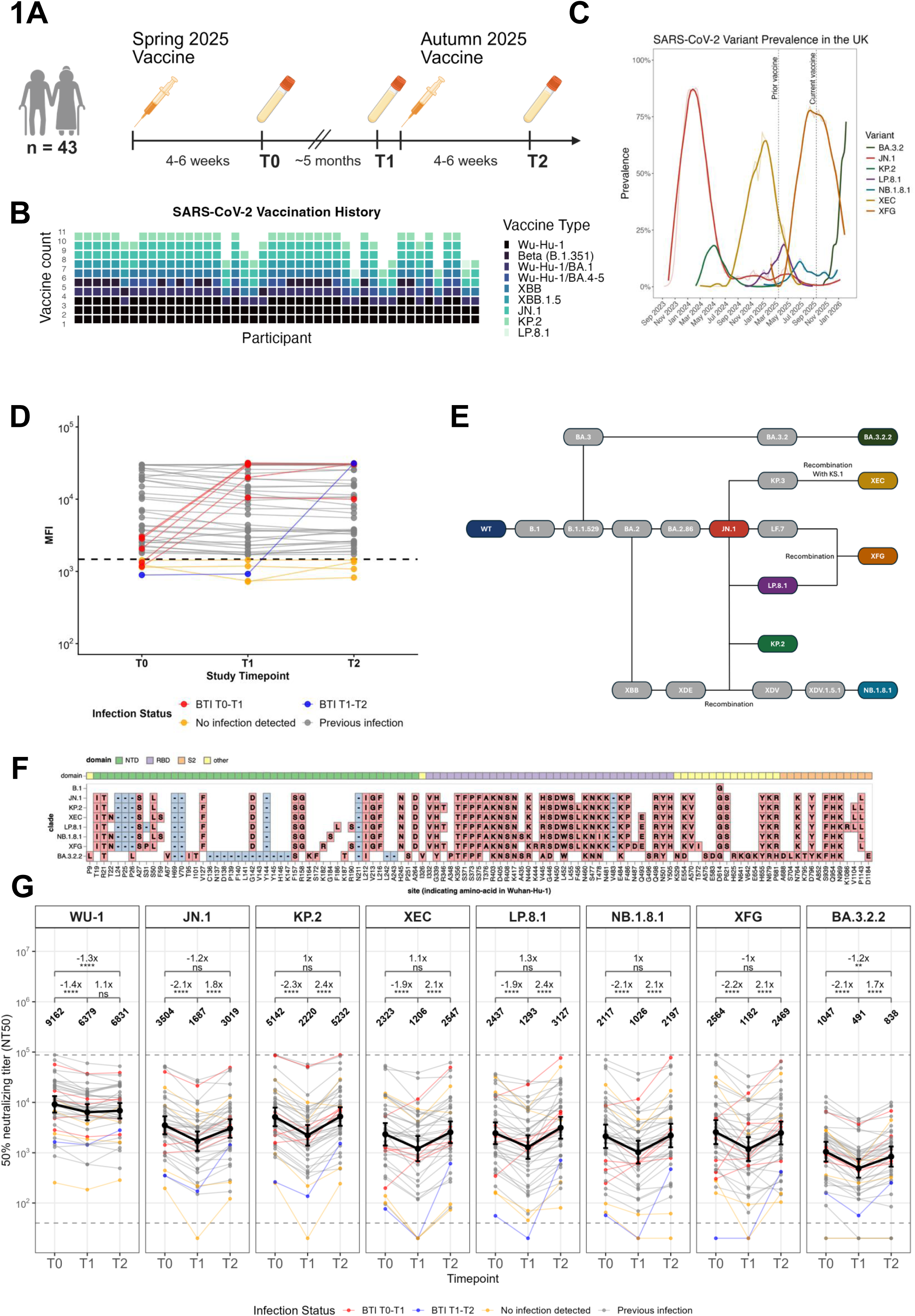
(A) Cohort schematic of individuals with two most recent doses. For 39/43 individuals in the cohort, a JN.1 vaccine was received in Spring 2025 and a KP.2 vaccine was received in Autumn 2025. T0, T1 and T2 indicate sampling timepoints that are referenced in this study. Created in BioRender. Morse, R. (2026) https://BioRender.com/sm9q7ti (B) Schematic of SARS-CoV-2 vaccination history for all participants included in this study. Blocks are coloured according to vaccine strain and ordered in the sequence participant received. Median number of 11 (IQR: 1) SARS-CoV-2 vaccinations from 2020-2025. (C) Prevalence plot indicating circulation of all SARS-CoV-2 variants included in the pseudotyped virus neutralisation assays in this study in the UK from September 2023-March 2026. (D) Plot of anti-nucleocapsid IgG as measured by Luminex binding assay. Five individuals were classified as having had a SARS-CoV-2 breakthrough infection between T0-T1, while one participant had a BTI between T1-T2. Four individuals fell below the assay cutoff for any prior SARS-CoV-2 infection, while the remainder (n = 33) are classified as having had at least one prior infection. (E) Schematic depicting the phylogenetic relationship between SARS-CoV-2 variants included in this study (coloured boxes). (F) Spike mutations present in variants included in this study. Amino acid insertions not depicted in this alignment. (G) Pseudovirus (PV) neutralisation of n = 43 individuals at ∼1 month post-JN1 dose (T0), ∼5-6 months post-JN1 dose and pre-KP2 dose (T1), and ∼1 month post-KP2 dose (T2). GMTs shown above each timepoint. Dotted lines indicate the lower (40) and upper (87480) limits of detection. Any values below the LLOD were set to half the LLOD.

**Table 1:** Cohort characteristics, vaccination histories, and study timepoints.

| Characteristic | N = 43<sup>†</sup> |
| --- | --- |
| **Demographics & Vaccination History** |  |
| Age, years — median (IQR) | 85 (75–88) |
| Female sex — no. (%) | 17 (40%) |
| Primary vaccine series |  |
| BNT162b2 | 27 (63%) |
| ChAdOx1 nCoV-19 | 16 (37%) |
| Total SARS-CoV-2 vaccine doses received — no. (%) |  |
| 7 | 3 (7.0%) |
| 8 | 6 (14%) |
| 9 | 1 (2.3%) |
| 10 | 6 (14%) |
| 11 | 27 (63%) |
| **Time between events, days — median (IQR)** |  |
| Spring vaccination to post-vaccination peak sample (T0) | 35 (33–40) |
| Post-vaccination peak sample to pre-autumn vaccination sample (T0-T1) | 133 (121–147) |
| Pre-autumn vaccination sample (T1) to autumn vaccination | 12 (6–18) |
| Autumn vaccination to post-vaccination peak sample (T2) | 33 (31–36) |
| Spring to autumn vaccination interval | 182 (174–190) |
| <sup>†</sup> Median (Q1–Q3); n (%) |  |

For the 2021 primary two-dose series targeting the original Wuhan-Hu-1 (Wu-1) spike, n = 27 participants (median age = 87, IQR 85-89, 37% female) received Comirnaty BNT162b2 (Pfizer-BioNTech) mRNA vaccination as their primary vaccine series, while n = 16 participants (median age = 75, IQR 71-79, 44% female) received the adenovirus-based AZD1222/ChAdOx1 nCoV-19 (AstraZeneca-Oxford) vaccine (Table 1, Supplementary Table 1). The age difference between primary series was expected given that BNT162b2 became available first and was offered to care home residents and individuals over age 80 who were deemed at highest risk. As variants began to circulate and evade immunity, booster vaccinations were recommended. By autumn 2025, participants had received a median of 11 (IQR 10-11) COVID-19 vaccine doses, including the 2025 booster dose (Table 1, Figure 1B). Booster doses included original and variant-based vaccines produced by Pfizer-BioNTech (Comirnaty BNT162b2) and Moderna (mRNA-1273), as well as a Beta variant-based protein subunit vaccine produced by Sanofi/GSK (AS03-adjuvanted VidPrevtyn Beta) (Figure 1B, Supplementary Figure 1A). The most recent booster doses targeted previously circulating variants that drove, or were predicted to drive, large waves of infection (Figure 1C)(24).

### Hybrid immunity in UK older adults

To interrogate antibody kinetics across the most recent doses, we tested sera obtained before and 1 month after the autumn 2025 vaccination, as well as 1 month before the previous vaccination (most commonly in spring 2025) (Figure 1A). We first tested these individuals for previous SARS-CoV-2 infection using an IgG binding assay against SARS-CoV-2 nucleocapsid (N) protein (Figure 1D). To define prior infection, we used a previously defined seropositivity threshold of 1472.8 mean fluorescence intensity (MFI) based on the value three standard deviations above the mean of n = 93 pre-pandemic serum samples(39). Values below the threshold were considered to indicate no detectable infection, although previous infection cannot be excluded because anti-N titers are known to wane over time(40). Values exceeding this threshold were considered to indicate a previous infection, while a two-fold or greater increase in IgG anti-N titers indicated a likely recent infection. At T0, n = 36/43 (83.7%) of individuals had detectable IgG anti-N demonstrating previous infection. Five breakthrough infections occurred between T0 and T1, with two individuals seroconverting, and one breakthrough infection that resulted in seroconversion also occurred within one month after the autumn 2025 vaccination (T1-T2) (Figure 1D).

### Kinetics of antibody responses to full-length spike after recent JN.1- and KP.2-based COVID-19 vaccines

We next aimed to assess how recent vaccination series contributed to antibody breadth and protection against circulating variants. We selected seven globally relevant variants that also circulated in the UK between September 2023 and March 2026, with Wu-1 D614G as a comparison (Figure 1C, E). Antigenic drift is evident between JN.1-derived variants, as compared to the pre-Omicron Wu-1 D614G and Omicron BA.3.2.2 derived from the saltation variant BA.3.2 (Figure 1F). One month after the previous dose (T0) received in autumn 2024 or spring 2025, neutralizing antibody titers remained highest against Wu-1 (Geometric mean titer, GMT: 9162), with 1.2- to 6.1-fold higher titers compared to the seven tested variants (Figure 1G, Supplementary Figure 1B-D). High titers against JN.1 (3504) and KP.2 (5142) were observed, as well as intermediate titers against XEC, LP.8.1, NB.1.8.1, and XFG (2323, 2437, 2117, and 2564, respectively). As expected due to significant antigenic drift and the beginning of circulation during the latter half of the autumn 2025 vaccination period, titers against BA.3.2.2 were significantly lower yet still reasonable (1047) (Figure 1C, F). Despite n = 39/43 (90.7%) individuals receiving two JN.1 doses in autumn 2024 and spring 2025, KP.2 titers were numerically higher ∼1 month after the second JN.1 dose (T0, KP.2 GMT = 5142, JN.1 GMT = 3504), suggesting good cross-protection conferred by the JN.1 vaccine against closely related variants with few amino acid substitutions (Figure 1G, Supplementary Figure 1B-D).

Neutralizing antibody titers against Wu-1 and all variants tested waned between doses, with Wu-1 waning the least (1.4-fold) and the seven variants waning by 1.9- to 2.3-fold (Figure 1G, Supplementary Figure 1D). However, one month after the most recent dose, neutralizing antibodies were significantly boosted for all variants tested by 1.7- to 2.4-fold (Figure 1G, Supplementary Figure 1D), indicating that booster vaccination increased antibody breadth after waning. There was no significant change in GMT between peak responses post-vaccination (T0 and T2) for JN.1, KP.2, XEC, LP.8.1, NB.1.8.1 and XFG, suggesting that an absolute titer ceiling may have been reached. Both Wu-1 and BA.3.2.2 titers, in contrast, significantly decreased between T0 and T2, indicating that the autumn 2025 KP.2 vaccination did not boost the antigenically distinct Wu-1 and BA.3.2.2 (Figure 1G, Supplementary Figure 1D). Taken together, these data suggest that booster vaccination increases neutralizing antibody titers and cross-neutralizing breadth against recently circulating variants, but that a threshold may be achieved for cross-neutralization without further targeted boosting.

### Hybrid immunity shapes antibody kinetics after JN.1-derived vaccination

To understand how hybrid immunity influences neutralizing antibody kinetics, we stratified the cohort by infection status. Individuals without recent infection or no detectable infection generally exhibited waning responses between doses that were boosted by the KP.2 vaccine except for Wu-1, whose titers continued to decline (Supplementary Figure 2A, yellow and gray). In contrast, the n = 5 individuals experiencing an infection between doses (T0-T1) demonstrated an absolute but nonsignificant overall decrease in Wu-1 titers, while showing increased titers for JN.1 lineage variants that surpassed the original titers at T0 (1.5- to 4.9-fold increase) (Supplementary Figure 2A, red). This suggests that infection drove titers beyond the ceiling achieved in individuals without recent infection, and that B cell responses were directed against new epitopes rather than conserved epitopes shared with Wu-1, thereby mitigating imprinting. As expected, the single individual who experienced a breakthrough infection after recent vaccination (T1-T2) had increased titers across Wu-1 and all variants, indicating some ‘backboosting’ of Wu-1 responses. However, fold-changes against variants were much higher: a 2-fold increase against Wu-1 compared to between 4.5- to 35.4-fold for all variants, which is consistent with mitigation of imprinting (Supplementary Figure 2A, blue). When stratifying by increase in titers between doses (T0 to T2), individuals with recent BTI tended to exhibit an increase in titers from T0 to T2 across almost all variants tested. In contrast, individuals without recent infection had a more heterogeneous response, with some exhibiting no boost between T0 to T2 and others exhibiting a boost only against some of the recent variants (Supplementary Figure 2B).

### SVNT enables analysis of breadth of RBD response

To understand whether the immunity generated by vaccination targeted the receptor-binding domain (RBD), as is often the case with mRNA vaccines, or other parts of spike, we utilized a surrogate virus neutralization (SVNT) assay involving the RBDs of 20 SARS-CoV-2 variants and SARS-CoV-1 (Figure 2A) and compared these results to our full-length spike pseudovirus assay. The highest titers after the spring 2025 vaccination were against Wu-1 (GMT: 1705), Gamma (1968), BA.1 (1710), and BA.2 (1739), consistent with variants to which study participants had been previously vaccinated or exposed (Figure 2B). As expected, titers waned between doses and then increased following the autumn 2025 booster (Figure 2B, Supplementary Figure 3A). Similar to pseudovirus results, titers either remained stable or decreased slightly between peak responses post-vaccination (T0-T2) (Supplementary Figure 3A), suggesting that a threshold was reached. In contrast to pseudovirus results, neutralizing antibody titers against Wu-1 RBD were significantly boosted after the autumn 2025 dose (T1-T2) (Figure 2C), suggesting that imprinting against the RBD remained, but that other non-RBD targets compensated for this and led to overall mitigation of imprinting. In addition, KP.2 anti-RBD titers were numerically lower than anti-RBD titers against JN.1, contrasting with pseudovirus results and suggesting that the high KP.2 neutralizing response could be driven in part by antibodies targeting non-RBD epitopes (Figure 2C). Similar to PVNT results, stratification by infection revealed that individuals with recent BTI tended to show an increase in titers from T0-T2 across most variants tested, whereas those without recent infection generally showed maintenance or a decrease in titers (Supplementary Figure 3B).

**Figure 2.**
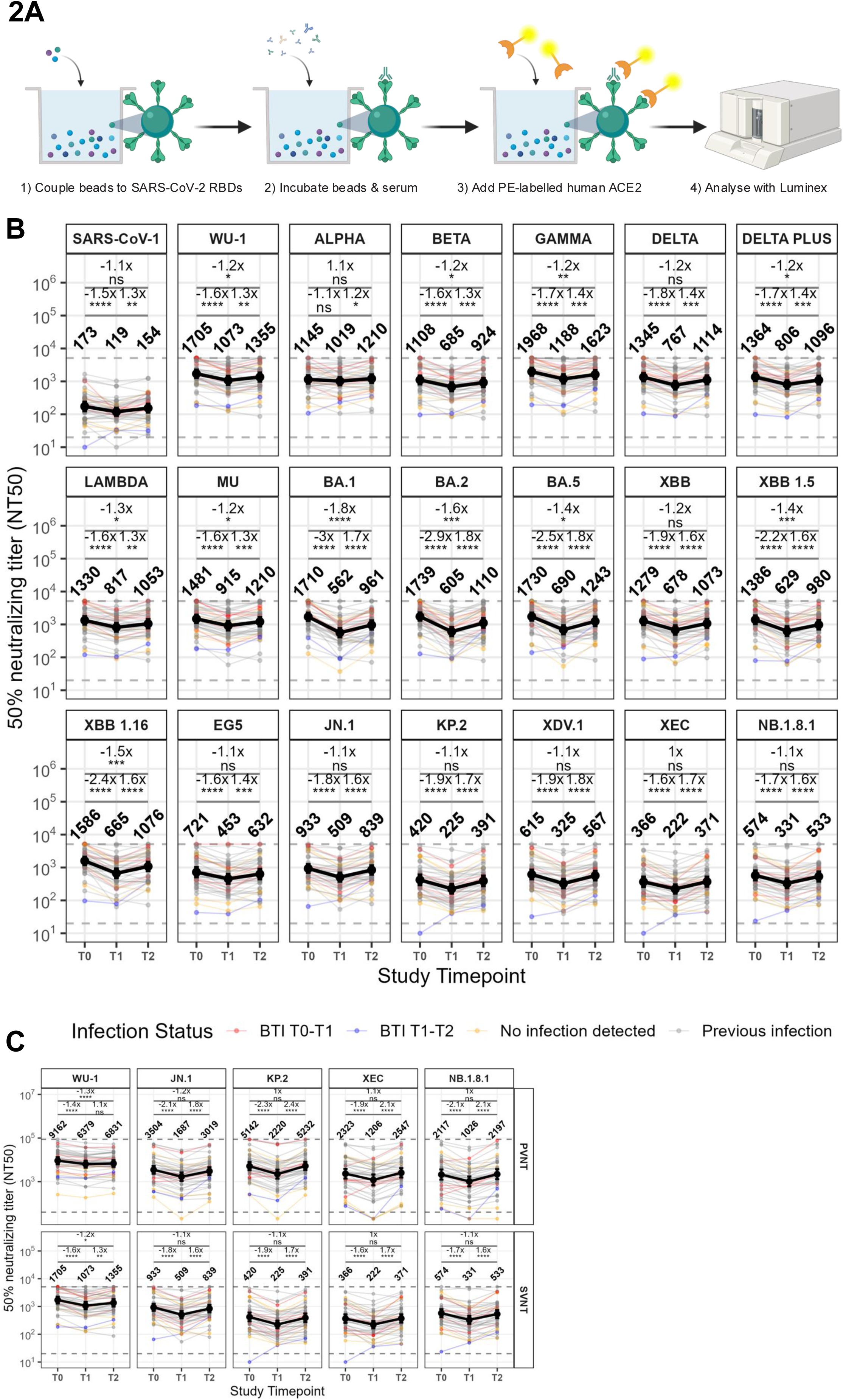
(A) Graphical schematic of Surrogate Virus Neutralisation Test (SVNT) assay performed using the Luminex system. MagPlex beads coupled to the RBDs of 21 variants were tested in a binding competition assay with ACE2 and detected by PE signal. (B) Neutralisation using SVNT assay of n = 43 individuals at ∼1 month post-JN1 dose (T0), ∼5-6 months post-JN1 dose and pre-KP2 dose (T1), and ∼1 month post-KP2 dose (T2). GMTs shown above each timepoint. Dotted lines indicate the lower (20) and upper (5120) limits of detection. Any values below the LLOD were set to half the LLOD. (C) Comparison of serum neutralisation by PV (top) and SVNT (bottom) assays for variants which were tested on both assays.

### Waning kinetics of variant-specific neutralizing antibodies in the elderly

Antibody waning was further examined by using a generalized linear mixed model that incorporated exposures from vaccination and BTI to estimate the variation in humoral response decay rate between variants. We first assessed serum neutralizing antibody half-lives over the three study timepoints via PVNT. We observed an antibody half-life of 179 days against Wu-1 full-length spike and significantly lower half-lives in the range of 43.7-70.1 days for the JN.1 lineage variants and 107 days for BA.3.2.2 (Figure 3A, bottom panel). Despite higher KP.2 neutralizing titers compared to JN.1 after both JN.1 and KP.2 vaccines (T0 and T2), the half-life of KP.2 neutralization (using antibodies against full-length spike) was much lower (47.6 days compared to 70.1 days), suggesting that a specific KP.2 boost may be needed to induce a durable variant-specific response (Figure 3A, bottom panel). Taken together, these data further suggest that multiple previous exposures to Wu-1 maintain long-term high absolute neutralizing antibody titers, whereas newer variant neutralizing titers with fewer exposures decay more rapidly.

**Figure 3.**
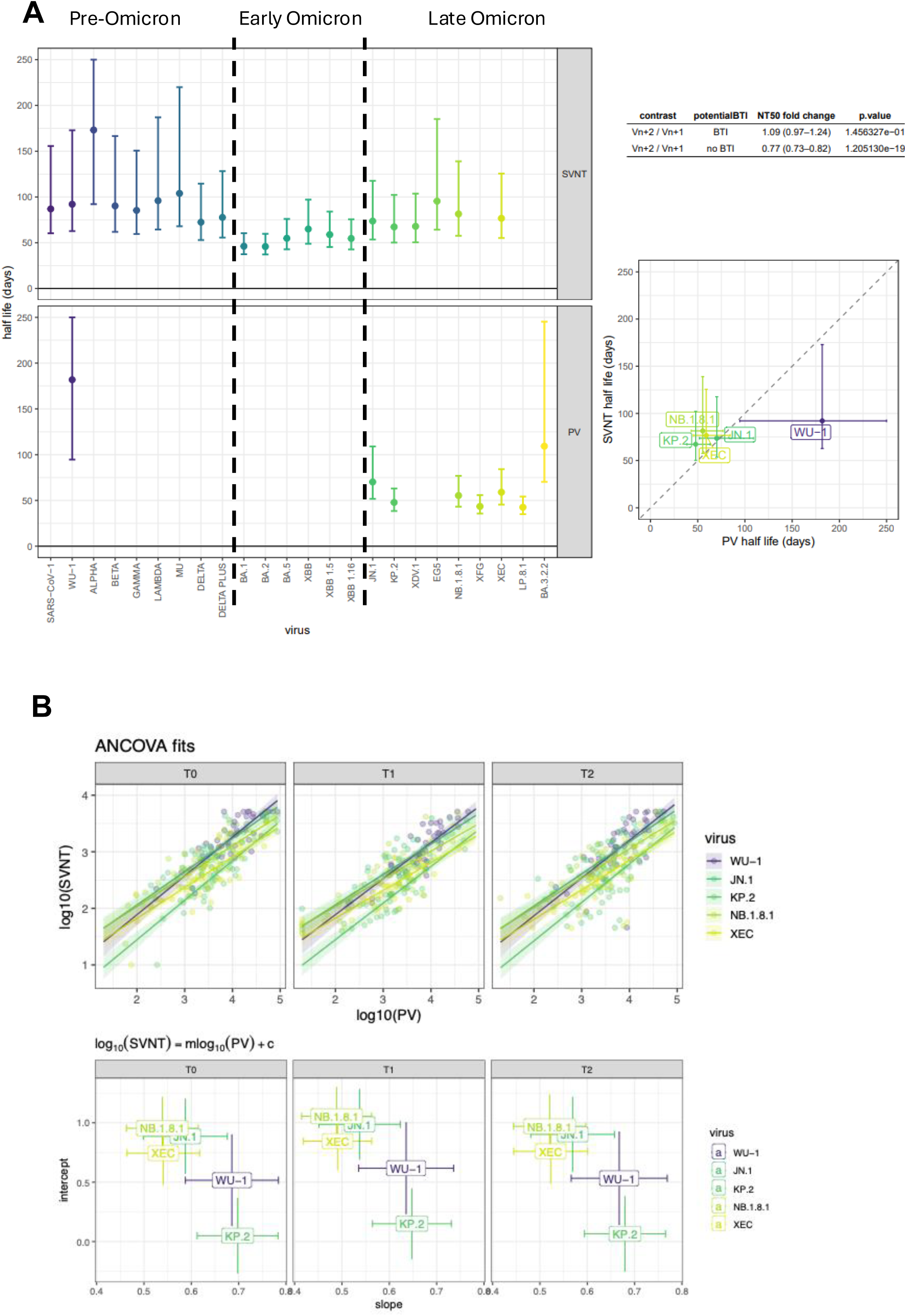
(A) Generalized linear mixed model for decay analysis. A generalised linear mixed model with a gamma distribution and a log link function was fitted for modelling the decay of antibodies. The derived half life and their 95% confidence interval are plotted, with upper limit of confidence interval capped at 250 days for ease of visualisation. (left) The modelled overall NT50 fold change between the Sp ring 2025 vaccine (Vn+1) and the Autum 2025 vaccine (Vn+2) based on if a breakthrough infection (BTI) happened between the vaccine. This shows plateauing of overall response that can be overcome by hybrid immunity (right upper). comparison of half -life and 95% confidence interval between variants where there are both PV and SVNT data. (right bottom) (B) ANCOVA fitted linear correlations between SVNT and PV assays, and the extracted intercept (c) and slope (m) coefficients of each overlapping variant (Wu-1, JN.1, KP.2, NB.1.8.1, XEC) across timepoint T0, T1, and T2.

We next assessed waning kinetics of antibodies targeting variant RBDs via SVNT. Whereas pre-Omicron RBD antibodies exhibited half-lives between 72-170.1 days (Wu-1: 91.3; Alpha: 170.1; all others: 72-102.9), earlier Omicron variants (BA.1 through XBB.1.16) decayed rapidly (45.7-64.7 days). However, decay rates stabilized for more recent post-XBB.1.16 variants (66.9-94.5 days), including JN.1 (73.2 days) and KP.2 (66.9 days), likely reflecting multiple recent exposures via JN.1 and KP.2 vaccinations or BTI (Figure 3A, top panel).

Given that the PVNT and SVNT assays use unique platforms to target different regions of the SARS-CoV-2 spike protein (full-length and RBD, respectively), we examined the relationship between the NT_50_s yielded by the two assays. We compared variants tested by both methods (Wu-1, JN.1, KP.2, NB.1.8.1, and XEC) by fitting a generalized linear model and using ANCOVA to isolate the effects of virus and timepoint. These data indicated that the virus variant, but not the timepoint at which neutralization was assessed, significantly changes the correlative relationship between the pseudovirus and SVNT NT_50_s (Figure 3B). Kinetic analyses confirmed this variant-specific divergence. In contrast to PVNT results, the half-life of KP.2 anti-RBD antibodies was similar, albeit slightly lower, compared to JN.1, suggesting that these RBD-directed antibodies decay at a similar rate but that KP.2 non-RBD neutralizing antibodies decay more quickly than JN.1, potentially due to fewer exposures (Figure 3A). We further observed a much shorter RBD-specific Wu-1 half-life compared to full-length spike (91.3 days vs. 179 days) (Figure 3A, right panel), suggesting that repeated hybrid immunity can assist in alleviating imprinting by shifting the RBD-specific humoral response such that ancestral RBD titers decay more quickly.

### Antibody landscapes in the hyper-vaccinated elderly

To visualize changes in the relationship between virus variants and neutralizing antibody responses, we constructed antigenic maps by translating fold-changes in neutralization titers towards different variants into antigenic map distances(12,41). The antigenic maps in Figures 4A and 5A show the antigenic relations between variants and sera across the three study timepoints. Viruses which cluster together on the map suggest a similar neutralization phenotype, while those that are distant indicate a distinct relationship between serum and antigen. By examining how the antigenic distance changes between timepoints, we take into account the change in titer against a variant in the greater context of other variants that share the antigenic map space, a nuance that is missed by the direct comparison of NT50s. In pseudovirus neutralization-derived maps depicting neutralization profiles to full-length spike (Figure 4A), JN.1 and viruses which are derived from this variant were clustered close to one another. Notably, BA.3.2.2 was antigenically distant to all other variants, demonstrating a distinct phenotype which is not mitigated by KP.2 booster vaccination. In PVNT-derived landscapes, there was a reduced neutralization profile across all variants from T0 to T1 (Figure 4B) as expected due to waning of antibody response over this 6-month window. However, at T2, after receiving the booster vaccine, we observe an appropriately skewed response that preferentially boosts the variants most similar to the vaccine strain (JN.1, KP.2, NB.1.8.1, XFG, XEC, LP.8.1). This is corroborated by significant increases in antigenic distance (Figure 4C) in sera towards these variants compared to Wu-1 between T0 to T1.

**Figure 4.**
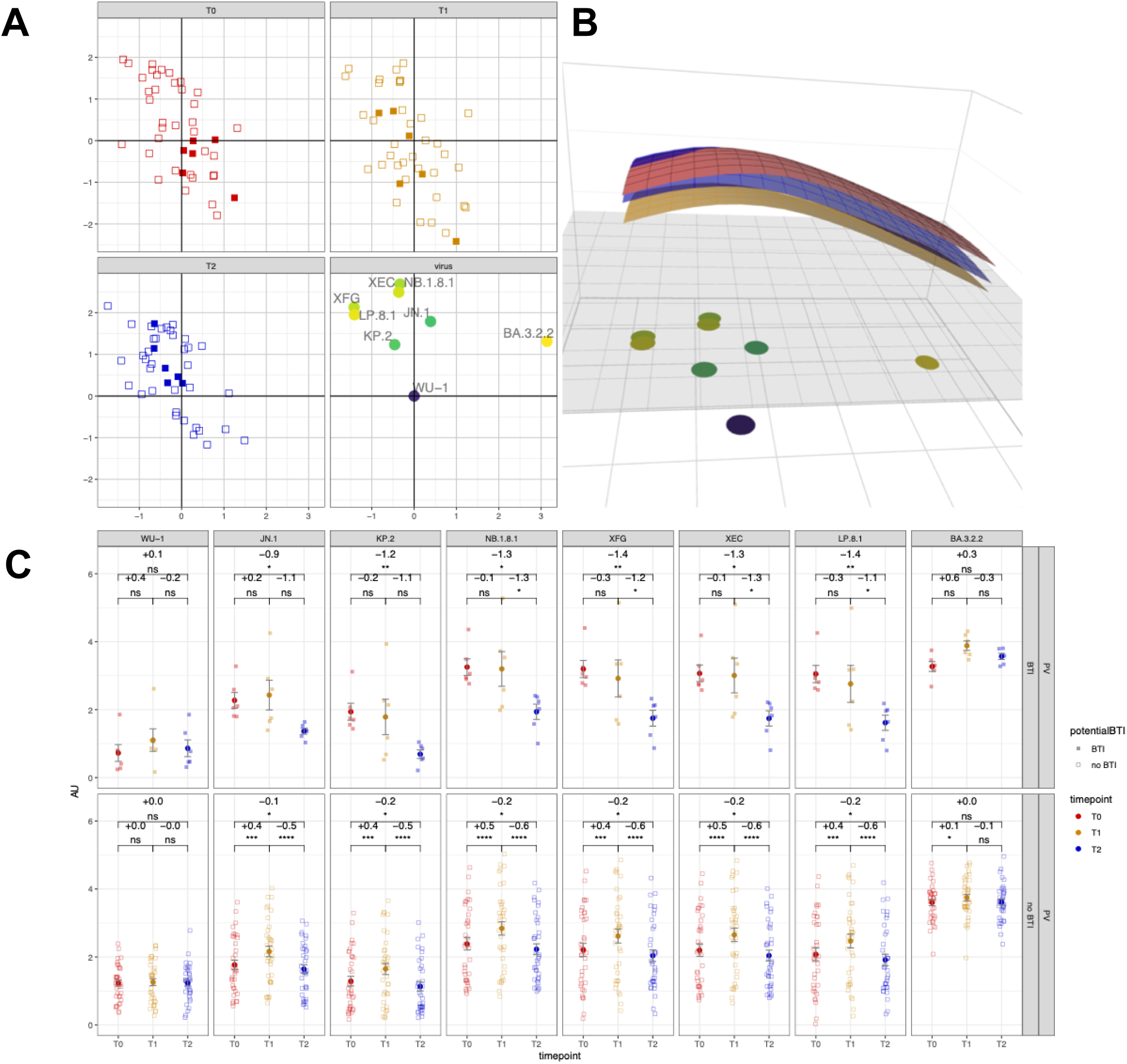
Deconstructed antigenic maps of pseudovirus neutralisation assay. (A) Deconstructed antigenic map split to depict serum from T0, T1, T2, and virus variant. One grid distance (antigenic unit, AU) in the map corresponds to a 2-fold difference in neutralizing titres. (B) Antibody landscapes for T0 (red), T1 (yellow), T2 (blue). The grey plane represents the lower limit of detection of the assay. The antibody landscape planes were fitted to the antigenic map in Figure 3A. (C) The antigenic distance in antigenic unit (AU) of each serum plotted against timepoint, facetted by the variant and the presen ce of breakthrough infection. The difference in the mean AU between each timepoint, as well as their statistical significance based on a two-tailed Wilcoxon test with Benjamini-Hochberg correction are annotated. * represents p<0.05, ** p<0.01, *** p<0.001, ns = not statistica lly significant.

**Figure 5.**
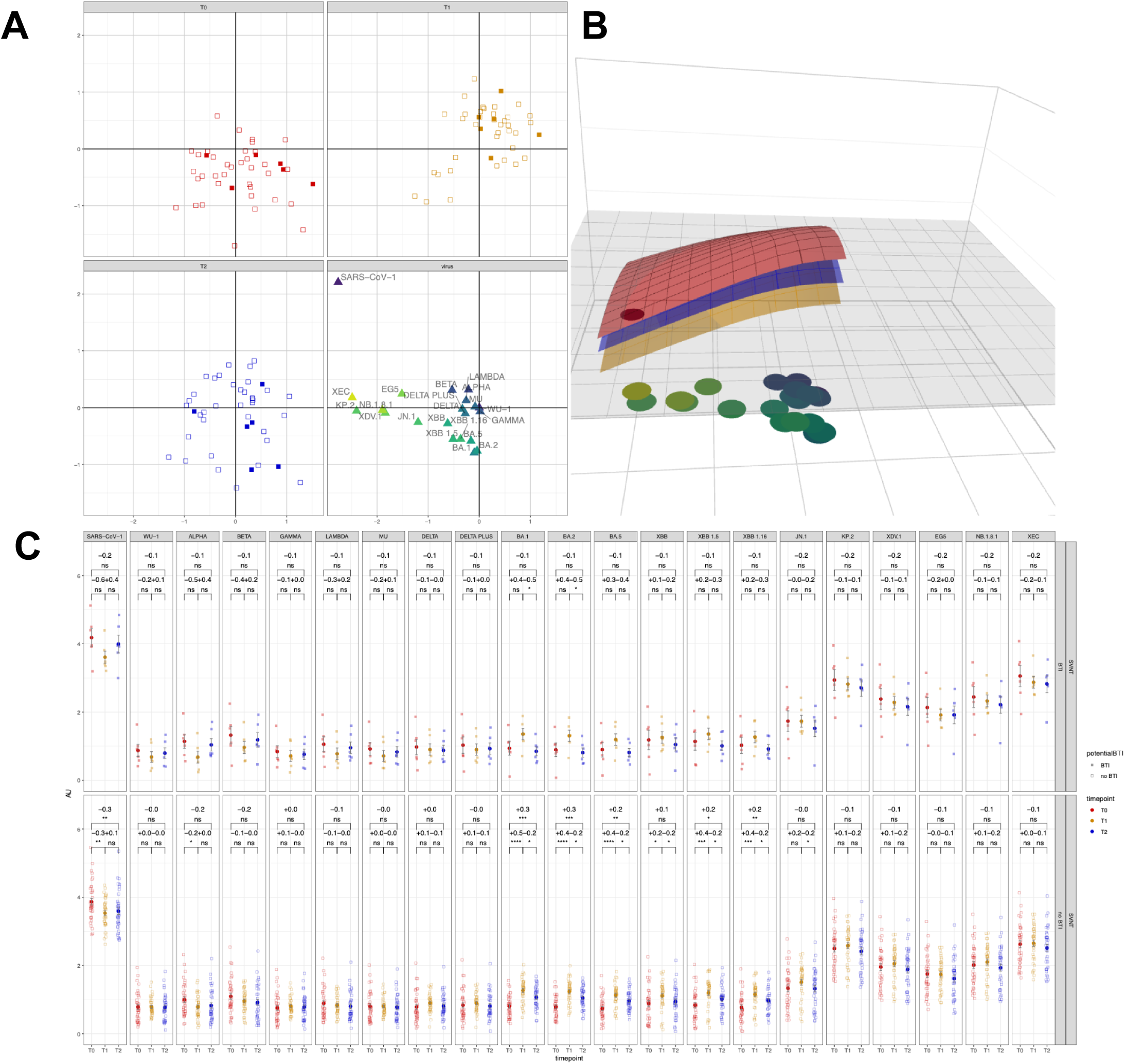
Deconstructed antigenic maps of surrogate virus neutralisation assay. (A) Deconstructed antigenic map split to depict serum from T0, T1, T2, and virus variant. One grid distance (antigenic unit, AU) in the map corresponds to a 2-fold difference in neutralizing titres. (B) Antibody landscapes for T0 (red), T1 (yellow), T2 (blue). The grey plane represents the lower limit of detection of the assay. The antibody landscape planes were fitted to the antigenic map in Figure 4A. (C) The antigenic distance in antigenic unit (AU) of each serum plotted against timepoint, facetted by the variant and the presen ce of breakthrough infection. The difference in the mean AU between each timepoint, as well as their statistical significance based on a two-tailed Wilcoxon test with Benjamini-Hochberg correction are annotated. * represents p<0.05, ** p<0.01, *** p<0.001, ns = not statistica lly significant.

SVNT-derived antigenic maps representing RBD-targeted neutralization demonstrate the relative antigenic distance of JN.1-derived RBDs compared to pre-JN.1 variants (Figure 5A). Like PVNT-derived landscapes, there was reduced neutralization across all variants from T0 to T1 (Figure 5B). The pattern of antigenic distances towards the RBD differ markedly to full-length spike, with KP.2, XDV.1, EG5, NB.1.8.1 and XEC being markedly distant to other variants (Figure 5A). These relatively poor RBD-centric neutralization responses to variants with similar RBDs, compared to similar full-length neutralization profiles, suggest that non-RBD antibodies are an important source of neutralizing breadth towards emerging variants. Notably, the boosting effect of the vaccine on RBD-directed neutralizing antibodies was substantially less pronounced compared to the effect seen in the PVNT assay (Figure 5C).

## Discussion

This study evaluated a UK cohort (n = 43) of highly vaccinated older adults (median age 85; median 11 vaccine doses) to map long-term, variant-specific immunity and neutralizing antibody responses elicited by JN.1-derived monovalent COVID-19 vaccinations. Following the UK spring 2025 vaccination round, we observed that neutralizing antibody titers waned across Wu-1 and all tested variants—JN.1, KP.2, NB.1.8.1, XFG, XEC, LP.8.1, and BA.3.2.2—over the ∼6 month period between doses, but were robustly boosted to similar peak levels after receiving an autumn 2025 vaccine. However, participants exhibited little neutralization against the antigenically distinct BA.3.2.2. Interestingly, although ancestral Wu-1 RBD neutralization followed conventional waning and boosting kinetics, neutralization against full-length Wu-1 spike steadily decreased across all timepoints tested, even in most individuals who experienced a breakthrough infection. This divergent pattern signals a progressive mitigation of immune imprinting with repeated Omicron exposures, suggesting that cumulative exposures via updated monovalent Omicron vaccination can reshape and redirect immunosenescent immune systems to broaden humoral immunity against emerging SARS-CoV-2 variants.

The antibody kinetics reveal preservation of peak immune responses but an accelerated waning timeline in older adults. Other studies examining contemporary monovalent vaccines targeting JN.1, KP.2 and LP.8.1 in younger and less frequently vaccinated cohorts demonstrate achievement of maximal absolute titers across most current variants, including JN.1, KP.2, LP.8.1, XEC, and NB.1.8.1. This boost appears to be short-lived as these titers continue to decay within 6-12 months post-vaccination (46) (47). Similarly, our elderly cohort achieved maximal titers across all variants tested except BA.3.2.2 following their autumn 2025 dose, with no significant difference between peak responses post-spring and -autumn 2025 dose (-1.2- to 1.3-fold change); this was generally observed for anti-RBD titers of the shared variants tested with SVNT (JN.1, KP.2, XEC and NB.1.8.1). However, the durability in our older cohort differs: a younger cohort receiving JN.1 and LP.8.1 vaccines one year apart had an average age of 33.8 (range 19-56) and significantly waning neutralizing antibody titers (1.7- to 3.3-fold decrease)(46). By contrast, we observed a similar 1.4- to 2.3-fold decrease in only 6 months between spring JN.1 and autumn KP.2 2025 vaccines. This comparable decay in just six months compared to one year aligns with reports that older adults above age 65 experience lower neutralizing titers and more pronounced waning compared to younger adults (29,42–45). Similarly, a recent study in older adult men that found a decline in vaccine efficacy—particularly against SARS-CoV-2 infection and hospitalization—beginning at 60 days post-vaccination through 120 days, although protection against death remained high(48). Together, these trends underline the shorter antibody half-lives in the Omicron era (45) and provide justification for bi-annual vaccination in a clinically relevant cohort of older adults.

Beyond temporal decay, our findings demonstrate a structural shift in the dominance hierarchy of the antibody response. Whereas the literature has predominantly focused on full-length spike neutralization, our comparison of pseudovirus data with the RBD-based SVNT reveals how updated variant matching alters immune targeting and displays distinct differences in RBD and full-length spike antibody kinetics. For instance, there was a notable divergence in decay rates: pre-Omicron and post-XBB.1.16 variants generally exhibited slower neutralizing anti-RBD titer decay compared to earlier Omicron lineages (BA.1 through XBB.1.16). This likely relates to exposure history, given the high number of Wu-1 exposures via Wu-1-based vaccination (3-4 monovalent, 2 bivalent doses) and JN.1 lineage doses (3) received in this cohort compared to only two XBB lineage doses. The ancestral Wu-1 response, however, revealed distinct differences between PVNT and SVNT. We observed significantly faster waning of Wu-1 anti-RBD titers compared to Wu-1 titers against full-length spike, suggesting that while repeated historical exposure to Wu-1 ensures long-term durability of absolute v against full-length spike, RBD-specific immunodominance wanes more quickly. Crucially, despite the significant post-KP.2 vaccine boost in Wu-1 anti-RBD, full-length Wu-1 neutralization continued to steadily decrease. These data indicate that the back-boosted, Wu-1 cross-reactive anti-RBD antibodies only constitute a small proportion of the overall neutralizing response and cannot compensate for the shift in immunodominance. Ultimately, these data suggest that despite extensive prior exposures to pre-Omicron variants via vaccination, multiple sequential exposures to updated JN.1-derived vaccines and infections may successfully reshape the immune hierarchy. This functional mitigation of imprinting occurs by redirecting or inducing *de novo* neutralizing responses to new epitopes both within and outside of the RBD.

This study further confirms that updated KP.2 and LP.8.1 vaccine campaigns increase neutralizing antibody breadth against JN.1-derived lineages, adding to our understanding of bi-annual vaccination patterns, but show that there is a ceiling reached when significant antigenic drift occurs. Similar to others, we find a limit in breadth when more divergent variants emerge with increased antigenic distance. LP.8.1 mRNA and protein vaccines have been shown to achieve high neutralizing titers similar to the most recent dose against more recent JN.1 lineage branches, such as XEC, LP.8.1 and NB.1.8.1, but fail to robustly cross-neutralize more antigenically distinct variants such as XFG and BA.3.2.2(46,49–51). In contrast, some found that LP.8.1 vaccines could shift immunodominance to also have high neutralization against XFG, but not BA.3.2(51,52). Our hyper-vaccinated cohort receiving a KP.2 vaccine, however, demonstrated maximal levels induced by JN.1-derived vaccination and at levels comparable to LP.8.1 and NB.1.8.1. This could suggest that the antigenic distance may be partially overcome by cumulative doses that broaden the memory B cell repertoire. Nevertheless, BA.3.2.2 remained modestly neutralized across age groups, geographies and infection histories, as neutralizing antibody titers remained lower than all other variants tested despite multi-dose vaccination of at least 5 doses(49–51,53,54). We note that this phenomenon may be largely due to the predominance of RBD in the down position in BA.3.2 variants, rather than simply alterations in epitopes recognized by neutralizing antibodies(55). Before the advent of monovalent Omicron vaccination, breadth of vaccination-induced responses did not extend to newer variants in this cohort such as XBB, BA.2.86 and related JN.1(32,56), demonstrating that boosters updated against more recently circulating variants are crucial for increasing neutralizing antibody breadth against very antigenically distinct variants.

A central finding was the mitigation of immune imprinting against the ancestral SARS-CoV-2 Wu-1 with repeated monovalent Omicron vaccination. The literature has shown divergent results regarding imprinting, a phenomenon that may be dependent on a cohort’s unique immune history. Some suggest that multiple breakthrough Omicron infection exposures can permit variant-specific neutralizing titers to meet or exceed levels against Wu-1 and drive *de novo* responses(19), although these findings were in a younger cohort with an inactivated primary series. In contrast, others found that updated mRNA variant-based vaccines along with BTI did not alleviate imprinting despite increasing breadth(20,57), though these cohorts were significantly smaller and younger than our cohort. Our data uniquely shows that continued repeated mRNA Omicron variant-based vaccination can alleviate immune imprinting in an older population, although our current data do not permit us to determine at what exposure number imprinting was overcome.

In younger adults receiving a KP.2 vaccine after 2-4 previous doses, KP.2 titers increased more than Wu-1 titers (4.1-fold vs. 1.5-fold, respectively), but Wu-1 titers remained highest, demonstrating that KP.2 vaccines improve breadth but that more doses are needed to overcome the back-boosting of Wu-1-specific immune memory(58). Similar patterns were observed in cohorts with 5-6 doses and JN.1 or KP.2 vaccine, where Wu-1 titers continued to increase by 2.3- and 3.9-fold, respectively(47). Importantly, our data disrupts this trend; although Wu-1 titers remain numerically higher due to cumulative exposures, the lack of Wu-1 neutralizing titer boosting after the autumn 2025 dose and the decline in full-length Wu-1 spike neutralization signals a shift in the memory B cell response to recall variant-specific memory B cells. This trajectory mirrors a recent sub-cohort of seven individuals receiving between 6-10 vaccine doses, where titers against JN.1-derived variants surpassed those of Wu-1 and Wu-1 titer fold changes were much lower than against variants(51).

Analysis of waning kinetics of neutralizing responses against a range of pre and post Omicron variants yielded interesting findings. The half-life of serum neutralization against Wu-1 (179 days) was substantially longer compared to JN.1 subvariants (43.7-70.1days), consistent with imprinted and longer-lived Wu-1-specific responses. Critically, BA.3.2.2 neutralisation was prolonged at 107 days compared to JN.1 subvariants, implicating imprinted ancestral variant targeting antibodies in neutralization of this saltation variant.

Overall, our data indicate that in the elderly, protective neutralising antibody responses against recent VOC can be achieved, with alleviation of immune imprinting by biannual, strain-updated booster vaccines. By investigating both RBD and full-length spike neutralization, we show that the alleviation of imprinting and preferential boosting of JN.1-derived antigenic targets is likely via non-RBD neutralizing antibodies. However, BA.3.2 reinforces the need to adapt vaccines to relevant variants in order to achieve maximal neutralization titers and protection from infection. With less frequent variant-specific boosting, imprinted responses against Wu-1 may re-gain dominance, as evidenced by analysis of longer serum Wu-1 neutralisation half-life. In the case of BA3.2.2 this imprinting may paradoxically confer relative protection against new saltation variants, given our data show slower waning of BA3.2.2 neutralisation compared to Omicron JN.1 lineages. Given imprinted antibodies against ancestral pre Omicron variants may still be relevant for protection against further highly immune evasive variants such as BA.3.2.2, therefore careful characterisation of population level COVID-19 humoral immunity is still warranted in order to plan updated vaccines.

## Limitations

Our sample size (n = 43) is modest and underpowered to detect smaller effects across multiple testing. Additionally, this cohort was recruited from a geographically distinct region in the UK with a unique virus and vaccine exposure history, thereby limiting generalizability. Our study lacked direct comparison to younger adults; however, older adults are the group currently prioritized for vaccination and many younger adults no longer receive COVID-19 vaccine boosters. Furthermore, we cannot definitively say that the KP.2 autumn 2025 vaccination alone reduced imprinting; this may have occurred at an earlier timepoint, and earlier pre- and post-vaccination samples require further assessment. Our study did not address T cell immunity, which is widely considered to represent broad and variant-agnostic protective responses that explain lower severity of infection in vaccinees who have breakthrough COVID-19. Given T cell escape is not thought to be common we did not explore this arm of immune response to vaccination in the elderly here. Finally, although this study examines the humoral component of immunity directed towards SARS-CoV-2, we did not characterize individual monoclonal antibodies in the repertoire for shifts in epitope targeting.

## Acknowledgments

We would like to thank all the study participants for their time and commitment to this research. We would also like to thank and acknowledge the efforts of the clinical and laboratory teams at the University of Cambridge for their assistance in participant recruitment, sample collection, and processing. We thank the Genotype-to-Phenotype National Virology Consortium (G2P-UK), and especially Dalan Bailey and Igor Santos, for providing SARS-CoV-2 spike-expressing plasmids. This work was supported by the Harding Distinguished Postgraduate Scholars Programme, the Gates-Cambridge Scholarship, the HKJC Global Health Institute, and Addenbrooke’s Charitable Trust. This work was also supported by the Cambridge University Hospitals Biomedical Research Centre.

## Author contributions

Conceptualization: R.B.M., D.J.S.E., M.T.K.C., R.K.G.; Methodology and Investigation: R.B.M., E.L.G, I.S., H.S, D.J.S.E., M.T.K.C., M.A., K.K., C.G., L.C.G., O.S; Data Curation and Analysis: R.B.M., D.J.S.E., M.T.K.C., R.K.G.; Funding Acquisition: R.K.G.; Resources: C.W.T., R.D., J.B., K.G.C.S., R.K.G.; Writing – original draft: R.B.M., D.J.S.E., M.T.K.C., R.K.G.

All authors approved the final version of the manuscript.

## Declaration of generative AI and AI-assisted technologies in the manuscript preparation process

During manuscript preparation, the authors used Gemini (Google) and Claude (Anthropic) to assist with language editing and coding support for data analysis. All scientific content and analyses were critically developed and verified by the authors. After using these tools, the authors reviewed and edited all AI-assisted outputs and take full responsibility for the integrity and accuracy of the manuscript.

## Materials and Methods

### Vaccine distribution in our cohort

In the UK, COVID-19 vaccines have consistently been offered to individuals over age 65 and to those with certain comorbidities such as immune compromised status. In our cohort, individuals received a primary two-dose series of either AstraZeneca, Moderna or Pfizer vaccines against the original Wu-1 strain, and a third Wu-1-based mRNA booster. After, vaccination histories diverged; some received a fourth monovalent dose, and some a bivalent original/BA.1 dose. These were followed by Original/BA.4-5 bivalent doses, and monovalent doses against XBB, XBB.1.5, JN.1, KP.2 and LP.8.1, respectively. Additionally, some individuals received a fifth or sixth Sanofi vaccine dose targeting the Beta variant.

We selected samples from all participants who had received a vaccine during the autumn 2025 COVID-19 vaccination campaign as well as a vaccine during either/both the autumn 2024 and spring 2025 COVID-19 vaccination campaigns (n = 43). At the first dose, the n = 43 participants had a median age of 80 years (IQR, 70-83). At the most recent sample, we estimated an additional five years added to each person’s age, providing a median age of 85 (IQR, 75-88). Participants were enrolled in the National Institute for Health Research (NIHR) BioResource (NBR) Centre study 118. Participants consented to providing relevant clinical data along with pre- and post-vaccination biological samples. The East of England Cambridge Central Research Ethics Committee (17/EE/0025) approved the study on April 28^th^, 2020.

### Binding antibody measurement

Binding IgG antibodies against SARS-CoV-2 nucleocapsid protein (N) were measured using the Luminex-based SARS-CoV-2-IgG assay by flow cytometry as previously detailed(40,59). Anti-N IgG cut-off was defined by ‘true’ positive convalescent sera and ‘true’ negative pre-pandemic sera, xzwhere the cut-off of 1472.8 MFI was three standard deviations above the ‘true’ negative mean(39).

### Cell lines

HeLa cells stably expressing ACE2 (HeLa-ACE2) and human embryonic kidney 293T (HEK293T) cells were cultured in Dulbecco’s Modified Eagle’s Medium (DMEM, Gibco), supplemented with 10% fetal bovine serum (FBS) and 1% penicillin/streptomycin. Cells were maintained in a humidified incubator kept at 37°C and 5% CO_2_. Cells were sub-cultured three times per week and kept below 25 passages.

### SARS-CoV-2 pseudotyped virus production and neutralization assays

In order to measure serum neutralizing antibodies to full-length spike of SARS-CoV-2 variants, lentiviral pseudotyped viruses were prepared by co-transfecting HEK293T cells with the following plasmids: HIV-1 Gag-Pol-Tat-Rev packaging plasmid p8.91, a firefly luciferase reporter gene construct with an HIV-1 packaging signal in the pCSFLW plasmids, and SARS-CoV-2 spike expression plasmids in the pcDNA3.1 vector. Spike expression plasmids encoded the spike protein of the following variants: Wu-1 D614G, JN.1, KP.2, XEC, LP.8.1, NB.1.8.1, XFG, BA.3.2.2, and included a deletion of the terminal 19 amino acids, as previously described(40,60,61). Pseudotyped virus (PV) stocks were titrated to ∼400,000 relative light units (RLU) on HeLa-ACE2 cells. PVs were incubated with pre-diluted participant sera for 1 hour at 37°C, following which this mixture was added to HeLa-ACE2 cells in white 96-wells plates, which were subsequently incubated for 48 hours at 37°C and 5% CO_2_. Following removal of supernatant, Bright-Glo (Promega) reagent was added to all wells and plates were read on a Promega Glomax Navigator Microplate Luminometer. Neutralization of PVs has been shown to correlate strongly with live virus neutralization(62). All neutralization assays were performed in technical duplicate and repeated across two independent experiments.

### Surrogate neutralization assay with panel of RBDs from variants

We performed multiplex SVNTs with AviTag-biotinylated SARS-CoV-2 RBD-conjugated MagPlex-Avidin beads (Luminex) and PE-conjugated human ACE-2 as previously described(63). In brief, heat-inactivated human serum samples at a starting dilution of 1:10 were titrated on a 96-well plate using a five point, four-fold dilution series in assay buffer (1% BSA + 1 mM NaCl in 1X PBS). SARS-CoV-2 RBD-coated beads (600 beads per antigen per well) were mixed and incubated with the serum dilution series for a final dilution of 1:20 on the plate’s top row. Sera and beads were incubated for 1 hour at 25°C shaking at 300 rpm. Next, 50 ul of 1000 ng/ml PE-conjugated ACE-2 was added to each well and incubated for 1 hour at 25°C shaking at 300 rpm. Finally, the plate was incubated on a magnetic plate holder for 2 minutes and washed twice with 1% BSA in 1X PBS before reading on the Bio-Plex 200 system (BioRad).

### Antigenic cartography

Antigenic maps were constructed using the antigenic cartography package Racmacs(64) (v1.2.9) in R(65) (v4.5.2) using pseudovirus and surrogate neutralization data. The table of neutralization titers is converted into a two-dimensional distance table by calculating log_2_ fold-change from the maximum titer for each serum to all other titers per serum. The co-ordinate map for each serum and variant is then optimized in 2000 steps. Euclidean distance in the map and their table distance is minimized. Maps were re-centered with Wu-1 being located on the origin. The mapDistances function was used to calculate antigenic distances. A detailed description of the method is given in the Racmacs package reference page, and in Smith et al(41,64). Antigenic cartography is based on multi-dimensional scaling, which is non-linear in nature, thus non-parametric statistical tests were performed using Rstatix v0.7.3(66) to compare antigenic distances between groups. Antigenic landscapes were projected and plotted using the antigenic landscape package ablandscapes v1.1.0(67). As previously described, for each serum group, single-cone landscapes were fit to each individual serum, and the geometric mean titer (GMT) landscape was calculated by taking the average of the landscape for the individual sera in that group(68,69).

### Definition of terms and statistical analysis

We defined vaccine breakthrough infection as a ≥2-fold increase in serum anti-N IgG between consecutive timepoints that exceeded the pre-defined cutoff for positivity based on previous studies (cite). Geometric Mean Titer (GMT) with standard deviation (SD) of neutralization antibody was calculated across time points. Fold-change (FC) was calculated by dividing the GMT of a given variant by the timepoint immediately prior and reported as mean FC and 95% CI. Baseline characteristics of participants were expressed as proportions and percentages for categorical variables and median inter-quartile range (IQR) for continuous variables. Two-tailed Wilcoxon signed-rank tests with Benjamini-Hochberg correction were used to compare neutralizing antibody titers between groups. Statistical analysis was performed using Rstatix v0.7.3.

### Generalized linear mixed model for decay analysis

A generalized linear mixed model with a gamma distribution and a log link function was fitted for modelling the decay of antibodies using the glmer function from lme4(70):

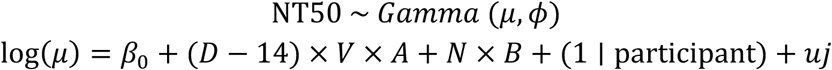

where D is the number of days since vaccination, V is the variant, A is the assay. To model the number of immunity exposures, N is the number of vaccines received by each participant, B is their breakthrough infection status. Individuals were treated as random effects to account for nonindependence by *uj* representing the participant-specific random intercept. Decay coefficients are converted to half-life measured in days for more intuitive comparison.

### Variation between pseudovirus and surrogate neutralization results

We fitted the following linear ANCOVA model using the lm function of stats, the R native package:

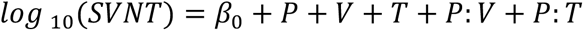

Where SVNT represents the surrogate neutralization assay NT_50_, P represents the log_10_(pseudovirus neutralization assay NT_50_), and the interacting factors represented by V (variant) and T (timepoint).

Next, analysis of covariance (ANCOVA) was performed using the package car v3.1-3(71) and coefficients extracted using the package broom v1.0.11(72).

---

**Supplementary Figure 1:**
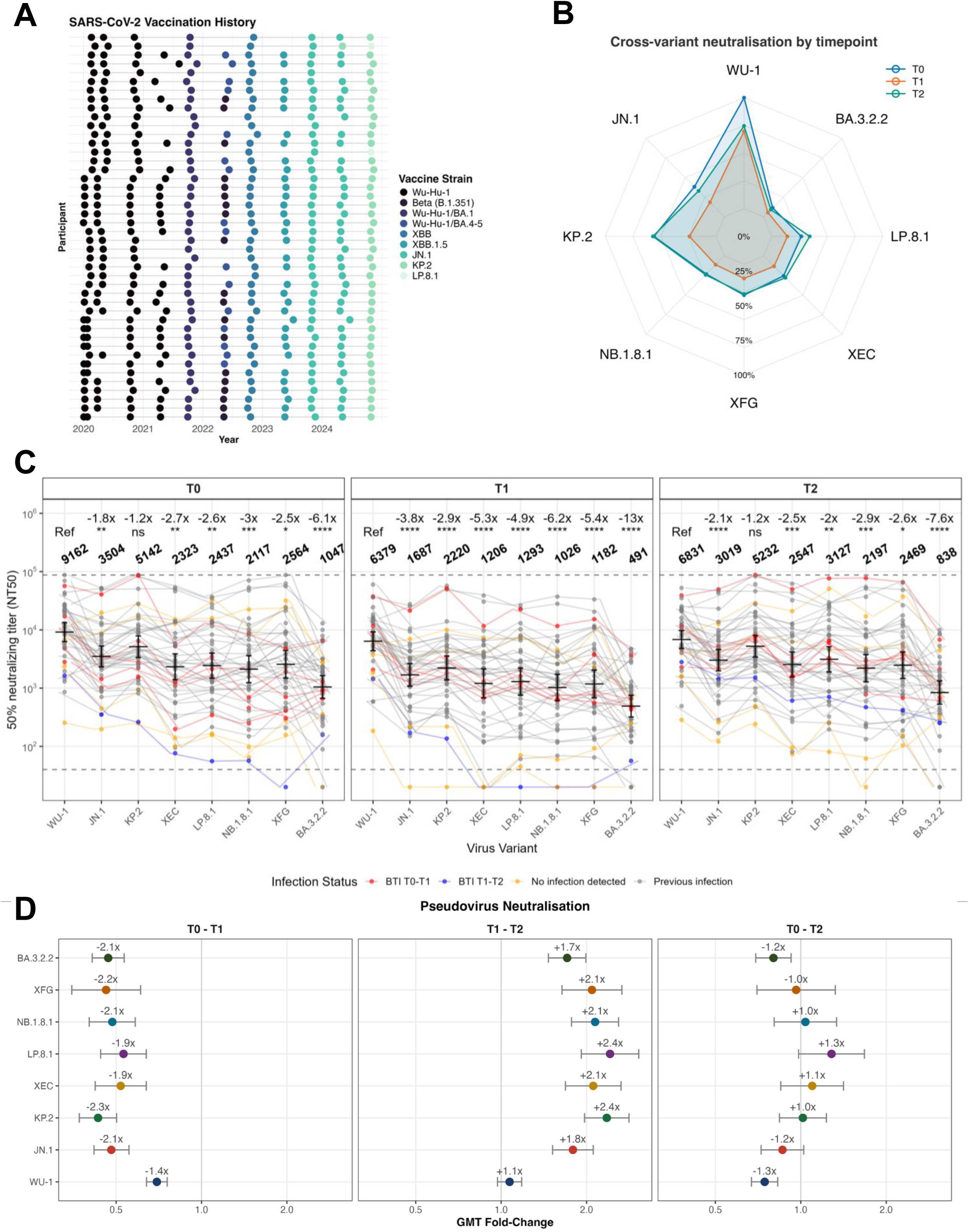
(A) SARS-CoV-2 vaccination history for all participants included in this study. Dots are coloured according to vaccine strain and indicate calendar date of vaccination. (B) Radar plot illustrating breadth of neutralising antibody responses across study timepoints. Data scaled relative to maximal response (Wu-1). (C) Neutralising antibody titres across study timepoints. Lines depict neutralising antibody titres for each individual across the variant panel over the three study timepoints. Grey lines indicate any previous SARS-CoV-2 infection, yellow denote no infection detected, blue lines indicate infection between T1-T2, and red indicate infection between T0-T1. Black bars indicate the GMT 95% confidence interval. The difference GMT between each variant relative to Wu-1, as well as their statistical significance based on a two-tailed Wilcoxon test with Benjamini-Hochberg correction are annotated. * represents p<0.05, ** p<0.01, *** p<0.001, ns = not statistically significant. (D) Strip plot of PV neutralising antibody GMT fold-change across study timepoints. Circles represent each variant and connected bars indicate the 95% confidence interval. Fold change of GMT neutralising titres between timepoints are indicated above each variant.

**Supplementary Figure 2:**
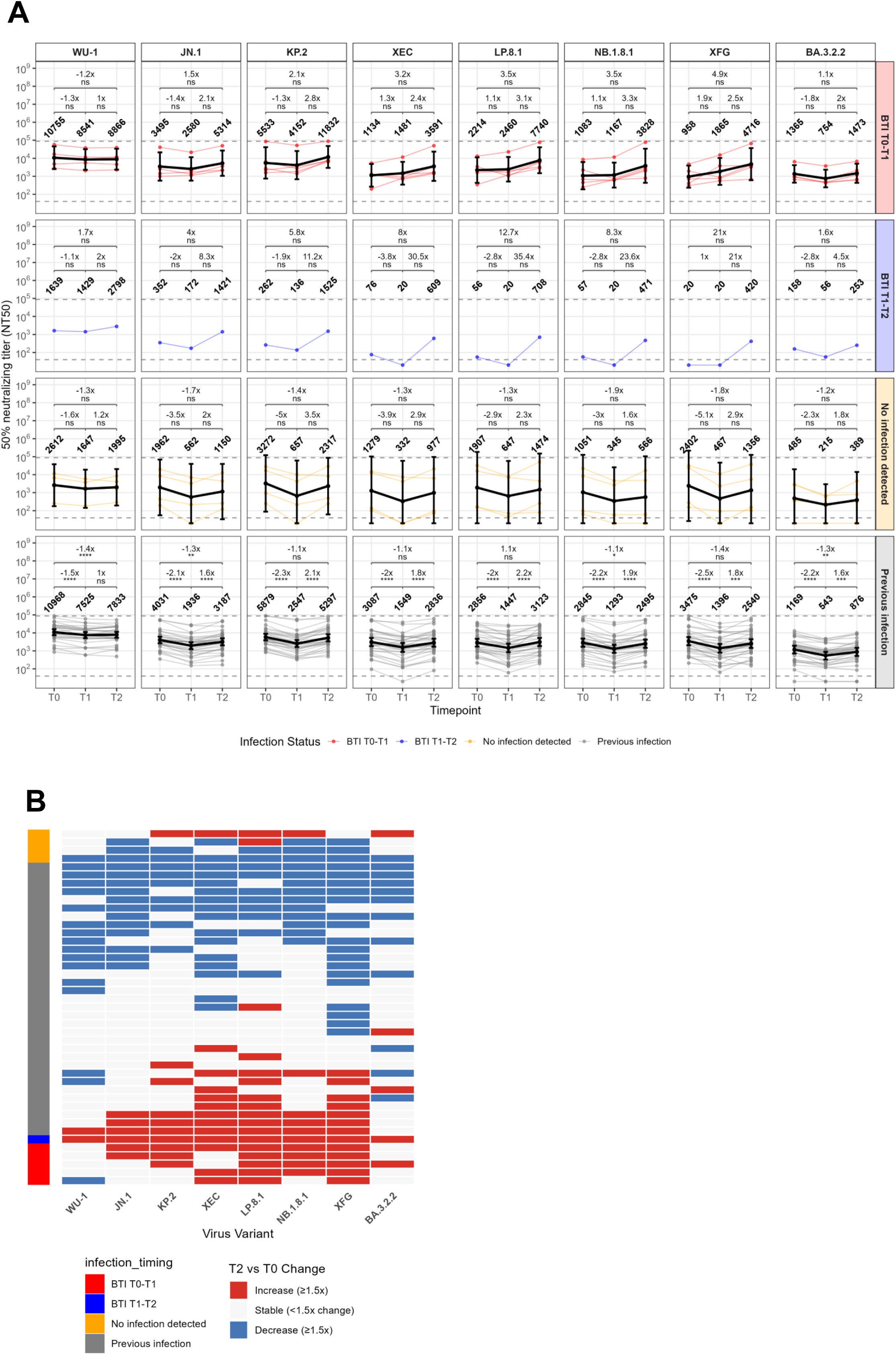
**A, B** Neutralising antibody titres across study timepoints stratified by N antibody status at baseline and during follow up time points. Lines depict neutralising antibody titres for each individual across the variant panel over the three study timepoints.

**Supplementary Figure 3:**
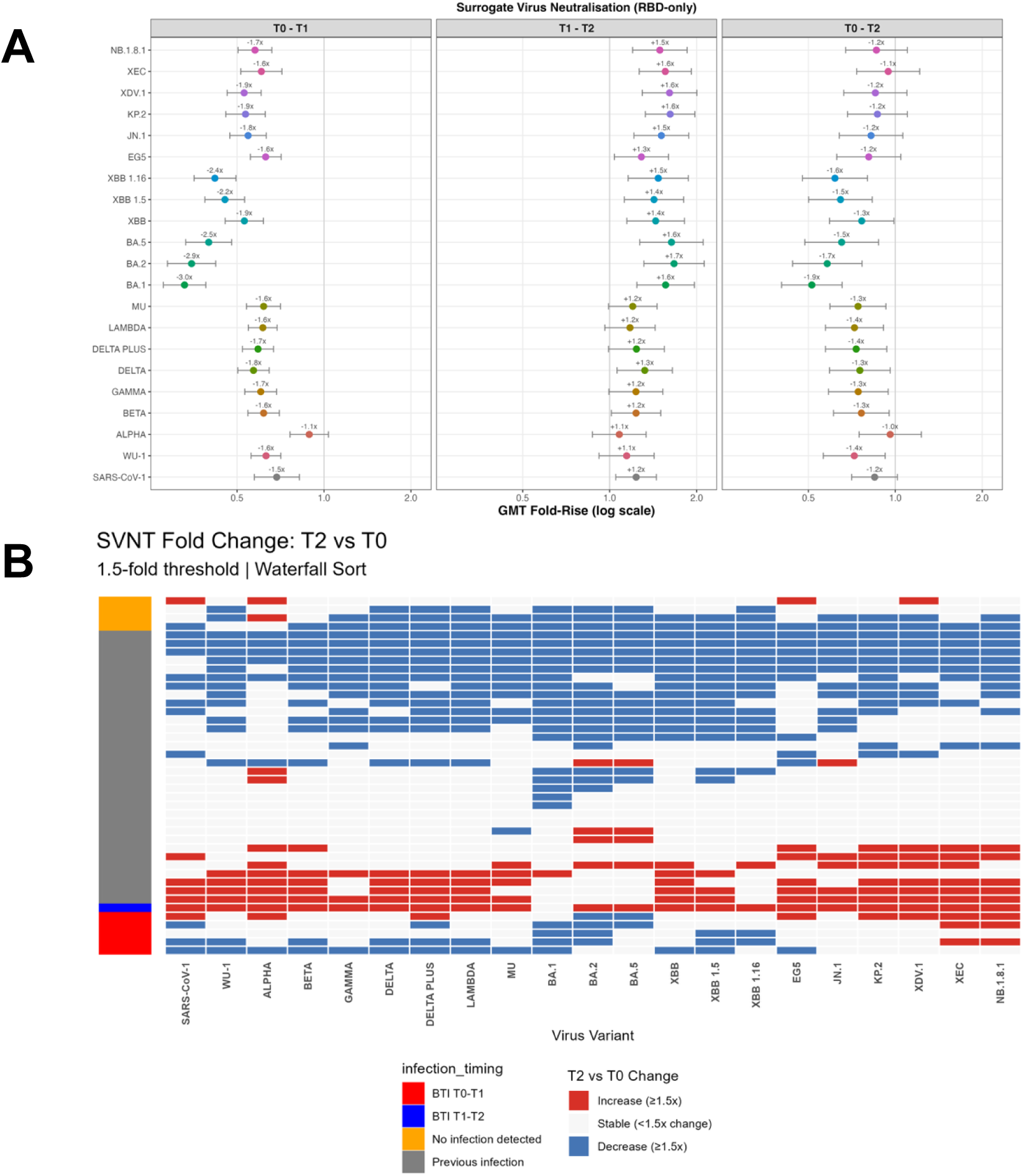
(A) Strip plot of SVNT neutralising antibody GMT fold-change across study timepoints. Circles represent each variant and connected bars indicate the 95% confidence interval. Fold change of GMT neutralising titres between timepoints are indicated above each variant. (B) Waterfall sort of SVNT GMT fold-change in neutralisation titres of study participants to 21 variants between T0-T2. GMT fold-change is indicated by colour of block for each individual for each variant. Blue fill indicates a >1.5 fold decrease in titre from T0-T2, red fill indicates >1.5 increase in GMT T0-T2, while light grey fill indicates a relatively stable titre with <1.5 fold-change. Strip along left of chart indicates SARS-CoV-2 infection status.

**Supplementary Table 1:** Cohort characteristics stratified by primary SARS-CoV-2 vaccine series.

| Characteristic | Overall<br>N = 43<sup>†</sup> | BNT162b2<br>N = 27<sup>†</sup> | ChAdOx1 nCoV-19<br>N = 16<sup>†</sup> |
| --- | --- | --- | --- |
| Demographics & Vaccination History |  |  |  |
| Age, years — median (IQR) | 85 (75–88) | 87 (85–89) | 75 (71–79) |
| Female sex — no. (%) | 17 (40%) | 10 (37%) | 7 (44%) |
| Total number of SARS-CoV-2 vaccine doses received — no. (%) |  |  |  |
| 7 | 3 (7.0%) | 0 (0%) | 3 (19%) |
| 8 | 6 (14%) | 3 (11%) | 3 (19%) |
| 9 | 1 (2.3%) | 0 (0%) | 1 (6.3%) |
| 10 | 6 (14%) | 4 (15%) | 2 (13%) |
| 11 | 27 (63%) | 20 (74%) | 7 (44%) |
| Time between events, days — median (IQR) |  |  |  |
| Spring vaccination to post-vaccination peak sample (T0) | 35 (33–40) | 34 (33–36) | 40 (35–43) |
| Post-vaccination peak sample to pre-autumn vaccination sample (T0-T1) | 133 (121–147) | 133 (119–145) | 136 (123–154) |
| Pre-autumn vaccination sample (T1) to autumn vaccination | 12 (6–18) | 12 (5–21) | 13 (8–16) |
| Autumn vaccination to post-vaccination peak sample (T2) | 33 (31–36) | 33 (30–36) | 34 (32–36) |
| Spring to autumn vaccination interval | 182 (174–190) | 176 (170–190) | 186 (179–194) |
| <sup>†</sup> Median (Q1–Q3); n (%) |  |  |  |

